# Dissection of protonation sites for antibacterial recognition and transport in QacA, a multi-drug efflux transporter

**DOI:** 10.1101/443523

**Authors:** Puja Majumder, Shashank Khare, Arunabh Athreya, Nazia Hussain, Ashutosh Gulati, Aravind Penmatsa

**Author notes:** Address for correspondence. Dr. Aravind Penmatsa, Assistant Professor, Molecular Biophysics Unit, Indian Institute Science, Bangalore, 560012 India., Email.

## Abstract

QacA is a drug:H^+^ antiporter (DHA2) with 14 transmembrane helices, that renders antibacterial resistance to methicillin-resistant *Staphylococcus aureus* (MRSA) strains, with homologues in other pathogenic organisms. It is a highly promiscuous antiporter, capable of H^+^- driven efflux of a wide array of cationic antibacterial compounds and dyes. Our study, using a homology model of QacA, reveals a group of six protonatable residues in its vestibule. Systematic mutagenesis resulted in identification of D34 (TM1), and a cluster of acidic residues in TM13 including E407 and D411 and D323 in TM10, as being crucial for substrate recognition and transport of monovalent and divalent cationic antibacterial compounds. The transport and binding properties of QacA and its mutants were explored using whole cells, inside-out vesicles, substrate-induced H^+^ release and microscale thermophoresis. We identify two sites, D34 and D411 as vital players in substrate recognition while E407 facilitates substrate efflux as a protonation site. We also observe that E407 plays a moonlighting role as a substrate recognition site for dequalinium transport. These observations rationalize the promiscuity of QacA for diverse substrates. The study unravels the role of acidic residues in QacA with implications for substrate recognition, promiscuity and processive transport in multidrug efflux transporters, related to QacA.

**Highlights:**

- A homology model of QacA with 14 TM helices was used to test the importance of acidic residues within the vestibule.
- Mono and Divalent cationic substrate recognition requires two sites D34 (TM1) and D411 (TM13).
- E407 (TM13) is important for protonation driven efflux.
- Substrate recognition of a divalent substrate, dequalinium occurs at E407 instead of D411 providing glimpses into the promiscuity of substrate recognition in QacA.

## INTRODUCTION

Efflux of antibacterial compounds is a major mechanism of acquiring multi-drug resistance in many pathogens [1, 2]. Efflux transporters of the major facilitator superfamily (MFS) are one of the largest groups of proteins involved in proton-coupled antiport in several pathogenic strains of Gram +ve and Gram -ve bacteria [3]. In methicillin-resistant *Staphylococcus aureus* (MRSA), chromosomally encoded antibiotic efflux pumps like NorA, NorB, NorC [4] and plasmid encoded antibacterial efflux pumps including QacA, QacB are involved in H^+^-driven transport [4, 5]. MFS transporters involved in antiport consists of candidates with 12 or 14 transmembrane (TM) helices, referred to as the DHA1 and DHA2 families, respectively [6]. Among DHA1 members, crystal structures are available for *E.coli* transporters MdfA, EmrD and YajR in multiple conformational states [7-9]. Extensive studies on MdfA and LmrP (*L. lactis*) revealed diverse aspects of their functional properties involving substrate promiscuity [10], H^+^:drug stoichiometry and sites for substrate recognition [11, 12]. Members of the DHA1 family also comprise chromaffin granule and synaptic vesicle monoamine transporters (VMATs 1 and 2) that share mechanistic similarities and transport serotonin, dopamine and noradrenaline into vesicles using H^+^-gradient [13, 14]. All DHA1 members previously studied have one or more acidic residues that are required for protonation and for lipophilic cation transport [12, 15, 16].

In contrast, with no available structures, the DHA2 family of antiporters have not been characterized extensively. QacA is a prototypical DHA2 member whose topology was determined to comprise 14 TM helices [17]. Molecules in the major facilitator superfamily have a conserved tepee-like architecture with two distinct six-TM helix bundles that are arranged with a pseudo two-fold symmetry [18]. Harnessing this property, QacA was modelled based on the distantly related 14 TM helix crystal structures of prokaryotic proton-dependent oligopeptide transporters (POTs) (~20% seq. identity) [19, 20]. POTs are also observed to bind peptides using an “aromatic clamp” that was proposed in MdtM, a DHA1 member [21, 22]. In the 14TM POTs, the two additional TM helices are observed as an insertion between symmetric helical bundles 1-6 and 7-12. As a result, TM helices 1-6 and 9-14 (in DHA2) have a pseudo two-fold symmetry that would allow rocking-switch movements to facilitate alternating-access on either side of the membrane [23].

Unlike MdfA, QacA is capable of H^+^: drug stoichiometries of 2 or greater [24] and QacA has an extensive substrate repertoire that includes nearly thirty organic monovalent and divalent cations, with antibacterial properties [25]. QacA is sensitive to inhibitors like verapamil and reserpine, of which the latter is a high affinity blocker of neurotransmitter transport, in both VMAT isoforms [24, 26]. A natural substitution of D323 to alanine is observed in QacB, a paralog of QacA with a corresponding inability to transport divalent substrates [17]. Antiporters require the protonation of at least one acidic residue to allow substrate binding and consequent H^+^-release, in the opposite direction to substrate flow [27]. However, it is frequently observed in transporters like LmrP [12], VMATs [28] and BbMAT [16], that more than one acidic residues exist within the transporter vestibule that can exchange two or more protons for every substrate molecule, rendering the process electrogenic [29].

Despite multiple studies on QacA’s prevalence, substrate repertoire and topology, experimental evidence that identifies residues critical for substrate recognition and protonation during transport process is lacking, presumptively due to the absence of a representative DHA2 structure. In this study, we employ a homology model of QacA to identify six acidic residues in the vestibule, which could play a role in protonation-coupled binding and transport. To investigate their role in the substrate recognition and transport of monovalent and divalent compounds, including ethidium (Et), tetraphenylphosphonium (TPP), pentamidine (Pm) and dequalinium (Dq), systematic mutagenesis of aspartate and glutamate residues in the vestibule of QacA was carried out. Efflux was monitored using whole cell-based and inside-out vesicle-based transport measurements to understand the effect of QacA mutants on substrate transport. Using purified QacA and its mutants, we further deduced the important roles of individual sites through substrate-induced proton release and direct-binding measurements using microscale thermophoresis (MST).

We identify that substrate recognition occurs at two distinct sites of the transporter, at D34 (TM 1) and D411 (TM 13), both of which are vital for binding and transport. Additionally, we observe that E407 is a likely protonation site that facilitates efflux. Rather interestingly, for a dicationic drug Dq, D411 ceases to be the second recognition site, whose role is replaced by E407, thereby providing some crucial insights into the promiscuous abilities of QacA to bind and transport diverse substrates using discrete sets of substrate recognition sites.

## RESULTS

### Homology model reveals six acidic residues in the vestibule of QacA

A 14TM homology model of QacA was built using prokaryotic POT structures [30] and MdfA as well as several other crystal structures as templates, using I-TASSER (Fig. 1a) [31]. QacA sequence lacking the region 432-474, corresponding to extracellular loop-7 (EL7), was used for the modeling study. The topology of QacA model, with the highest score, resembles POTs, where in TMs 1-6 and TMs 9-14 have an inherent pseudo-symmetry and TMs 7 and 8 form an insertion that connects the two domains that undergo alternating-access [23]. The boundaries of the modeled TM helices broadly adhere to the previously determined experimental topology [17]. The model exhibits an inward-open conformation with solvent accessibility towards the cytosolic compartment. We observed six acidic residues, lining the vestibule of QacA at different levels including D34 (TM1), D61 (TM2), D323 (TM10), E406 (TM13), E407 (TM13) and D411 (TM13) (Fig. 1a, b and S1). With the exception of D323 and E406, each of the acidic residues is well conserved among closely related QacA homologues. The accessibility of individual residues was verified by mutating each of the six residues to a cysteine. The absence of cysteine residues in the primary sequence of QacA facilitated our analysis of solvent accessibility to acidic residues within the vestibule, using PEG-Maleimide (PEG-Mal) that covalently binds to free thiol groups [32]. Single cysteine substitutions of QacA at E406, E407 and D411 exhibited greater propensity for PEG-Mal modification induced gel shifts, whereas substitutions at D34, D61 and D323 exhibited minimal labeling, suggesting reduced PEG-mal (and thus solvent) accessibility (Fig. 1c). The greater accessibility of PEG-Mal to residues towards the cytosolic part of the transporter vestibule allows us to infer that QacA likely exists in the inward-open conformation in detergent micelles and the homology model reflects this orientation.

**Figure 1.**
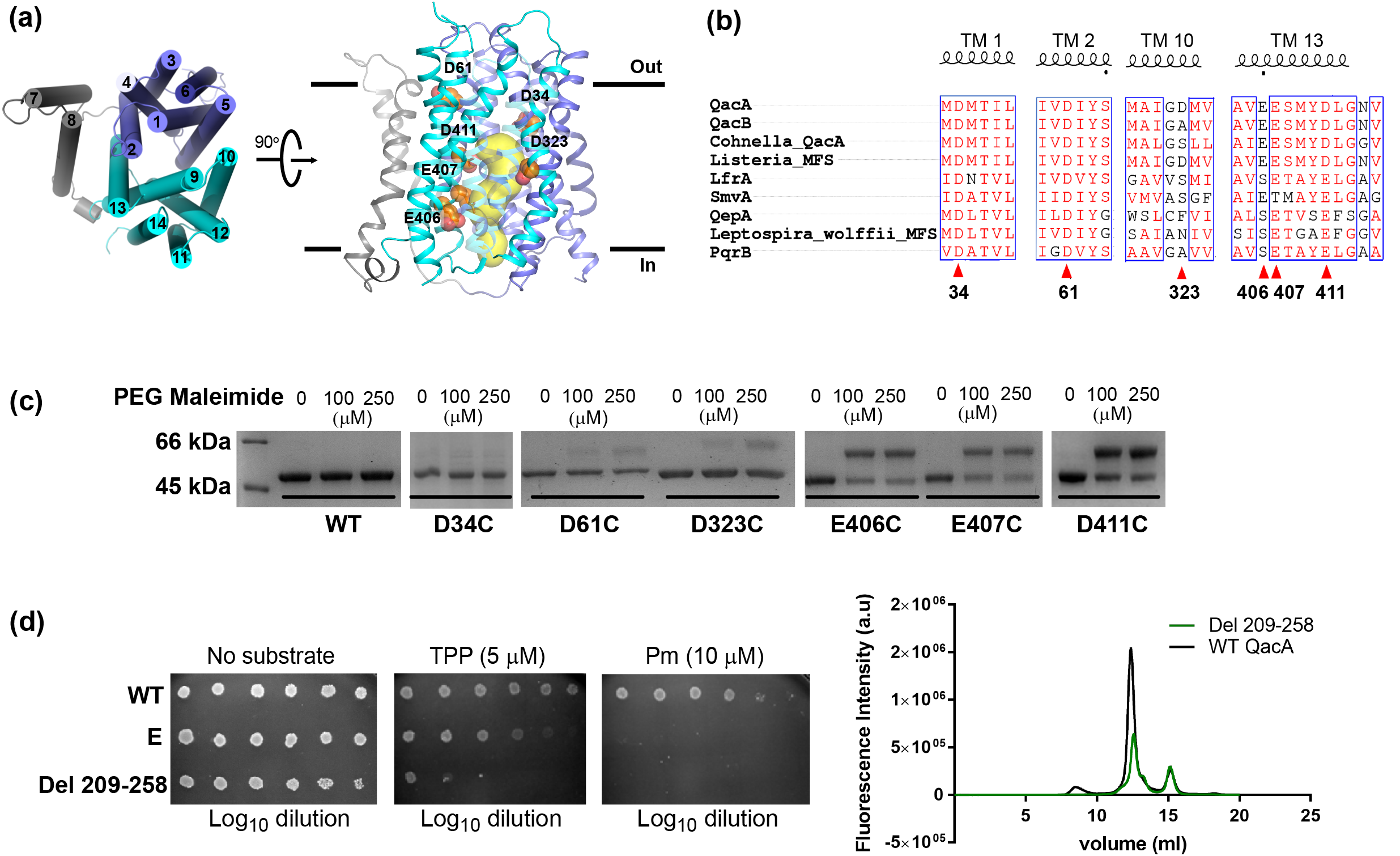
Homology model of QacA. (a) Model shows TMs 1-6 (dark blue) and TMs 9-14 (cyan) displaying pseudo two-fold symmetry. TMs 7 and 8 (grey) connect the two domains. Acidic residues in the vesti-bule are indicated in orange. The model displays an inward-open conformation with solvent accessible vestibule colored in yellow. (b) Multiple sequence alignment of QacA and its homologs across various species. Acidic residues mapped in QacA vestibule are pointed with red arrows. (c) PEG-maleimide accessibility assay indicates band shifts to higher molecular mass due to PEG-maleimide covalently modifying individual cysteine mutants. (d) The loss of activity for TM 7 and 8 deletion construct (Δ209-258) observed in presence of monovalent (TPP) and divalent (Pm) cations (left). FSEC profile of the deletion construct displays a homogenous protein peak (right).

We evaluated the model by exploring the functional importance of the region corresponding to the TM helices 7 and 8 by deleting a stretch of residues from 209-258. Despite the deletion, the protein expression displayed a homogenous profile as observed using fluorescence-detection size exclusion chromatography (FSEC), suggesting structural integrity (Fig. 1d). The deletion construct suffered a complete loss of function as observed using survival assays, in comparison with WT QacA. The observation suggests that TMs 7 and 8 in QacA, play a significant role in mediating efflux, in accordance with the observations with Tet(L) TM7 and 8 deletion that loses the ability to provide tetracycline resistance [33]. The ability of WT QacA to transport drugs was tested at high extracellular pH (pH 8), where the pH gradient is insignificant across *E. coli* membranes (Fig. 2). *E. coli* expressing WT QacA could robustly protect cells against TPP, Pm and efflux Et when the extracellular pH was 6 whereas this ability was compromised when the extracellular pH was enhanced to pH 8 suggesting that transport in QacA is driven primarily by pH gradient and to a lesser extent through membrane potential.

**Figure 2.**
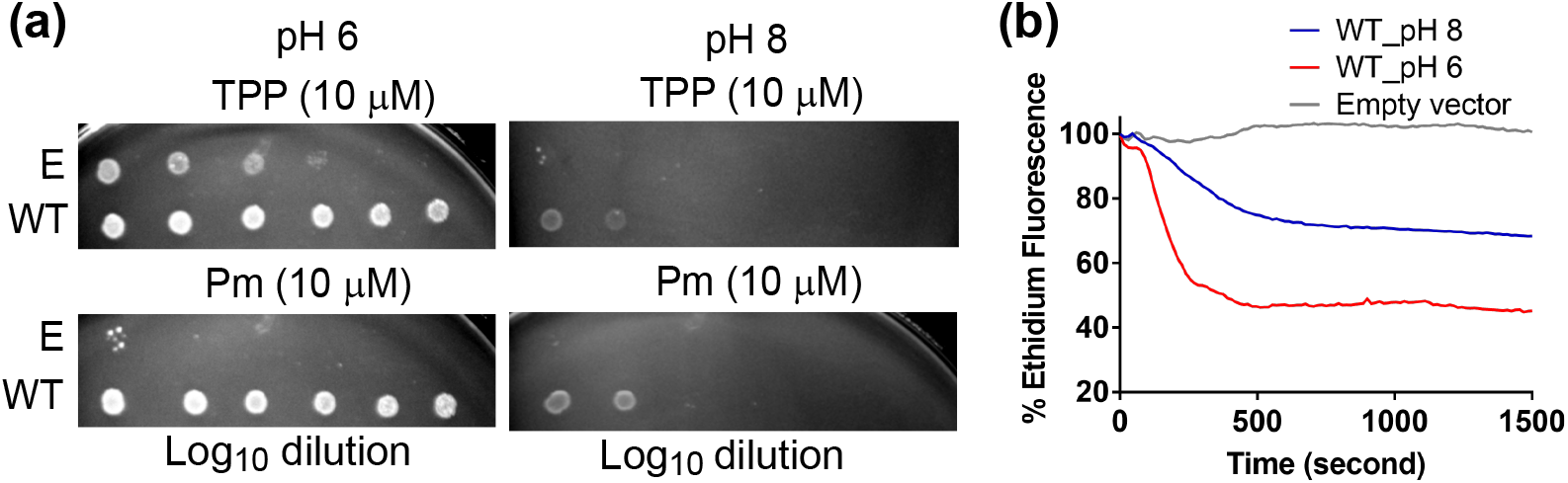
ΔpH dependence of QacA’s activity checked by (a) survival and (b) whole cell (ethidium) efflux assays done at external pH of 6 and 8. All assays were independently performed at least twice.

Proton-driven transport of positively charged or polar molecules through QacA requires acidic residues inside the vestibule that are capable of reversible protonation and deprotonation upon lipophilic cation binding. The significance of acidic residues observed within the QacA vestibule was characterized through individual residue substitutions to alanine and isosteric asparagine or glutamine. Constructs with and without a GFP-His_8_ tag at the C-terminus were built to facilitate expression analyses through FSEC.

### Behaviour of WT QacA

QacA was purified as described earlier [34] using affinity purification with a moderate yield and high level of purity (Fig. 3), using undecyl-β-D-maltopyranoside (UDM). Purified QacA displays a higher oligomeric species in solution whose molecular mass corresponds to a QacA trimer and heavy aggregation was observed when concentrated beyond 1 mg/ml. The propensity to aggregate also precluded structural and biophysical studies requiring concentrated samples of QacA. The binding and competition studies performed with purified QacA were therefore carried out in dilute samples where QacA displays a predominantly monomeric species. Before assessing the roles of individual residues in transporting the cationic substrates, WT QacA was assayed in whole cells, inside-out vesicles and as detergent isolates for its behavior in the same experiments. Et transport property was checked in whole cell-based efflux assay, where a relative decrease in the fluorescence of Et was correlated with its efflux from energized cells. To assess the same in case of TPP, Pm and Dq, inside-out vesicles were prepared and ACMA was used as a pH dependent fluorescent probe. Dequenching of ACMA fluorescence was used as a measure of substrate induced proton antiport (Fig. 4a). Binding of substrates to the purified QacA was evaluated using microscale thermophoresis that allowed rapid estimation of binding affinities (Fig. 4b). In substrate-induced proton release assays, unbuffered solution of purified QacA was acidified followed by a step-wise addition of fixed concentrations of the substrate [22]. The release of protons was monitored through quenching of a pH sensitive probe fluorescein (Fig. 4c, S5).

**Figure 3.**
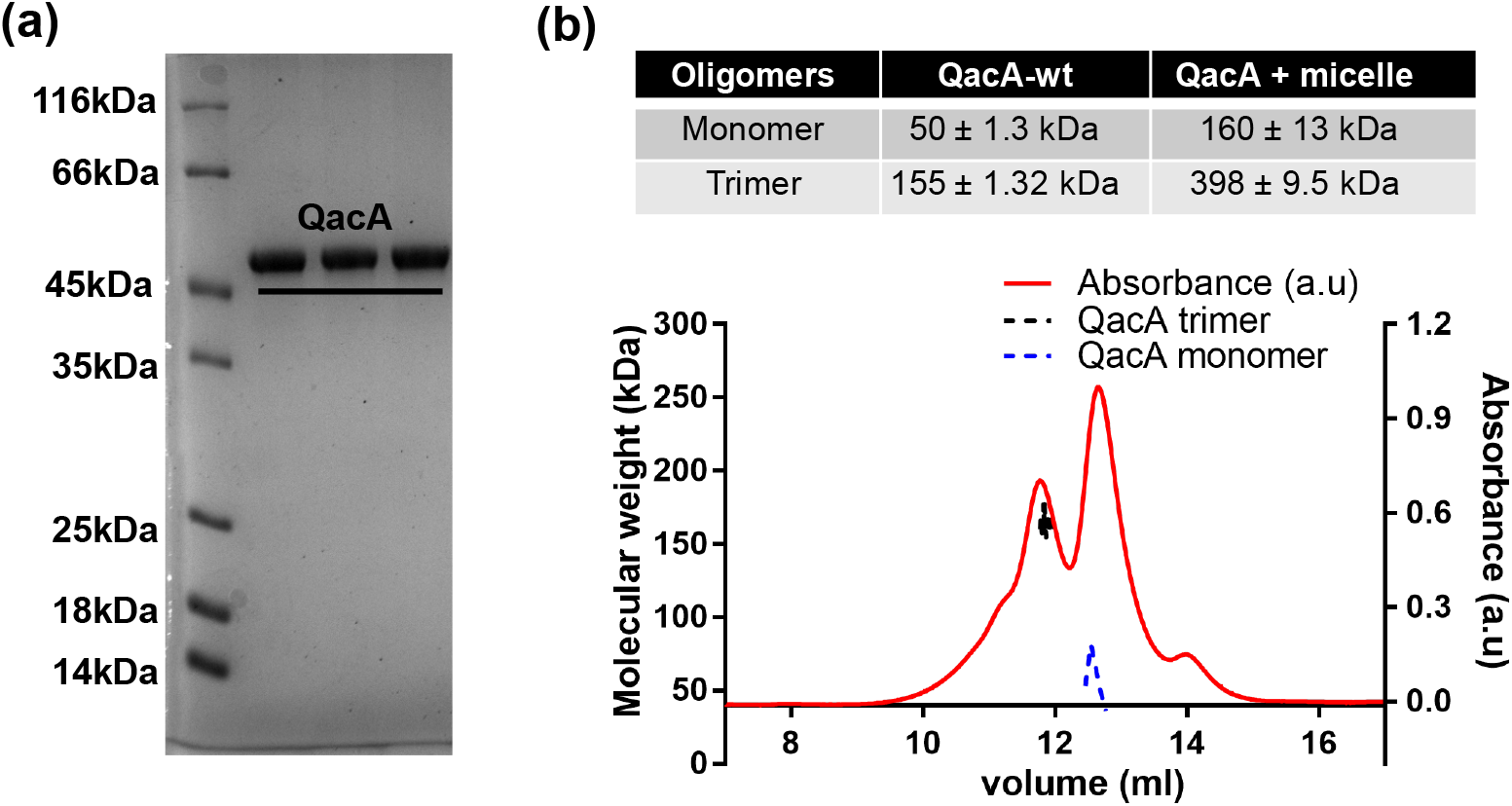
QacA purification profile in UDM (n-undecyl β-D-maltopyranoside) (a) Ni-NTA purified fractions of WT QacA in SDS-PAGE, (b) SEC-MALS profile of QacA in UDM to check the oligomeric status of the protein.

**Figure 4.**
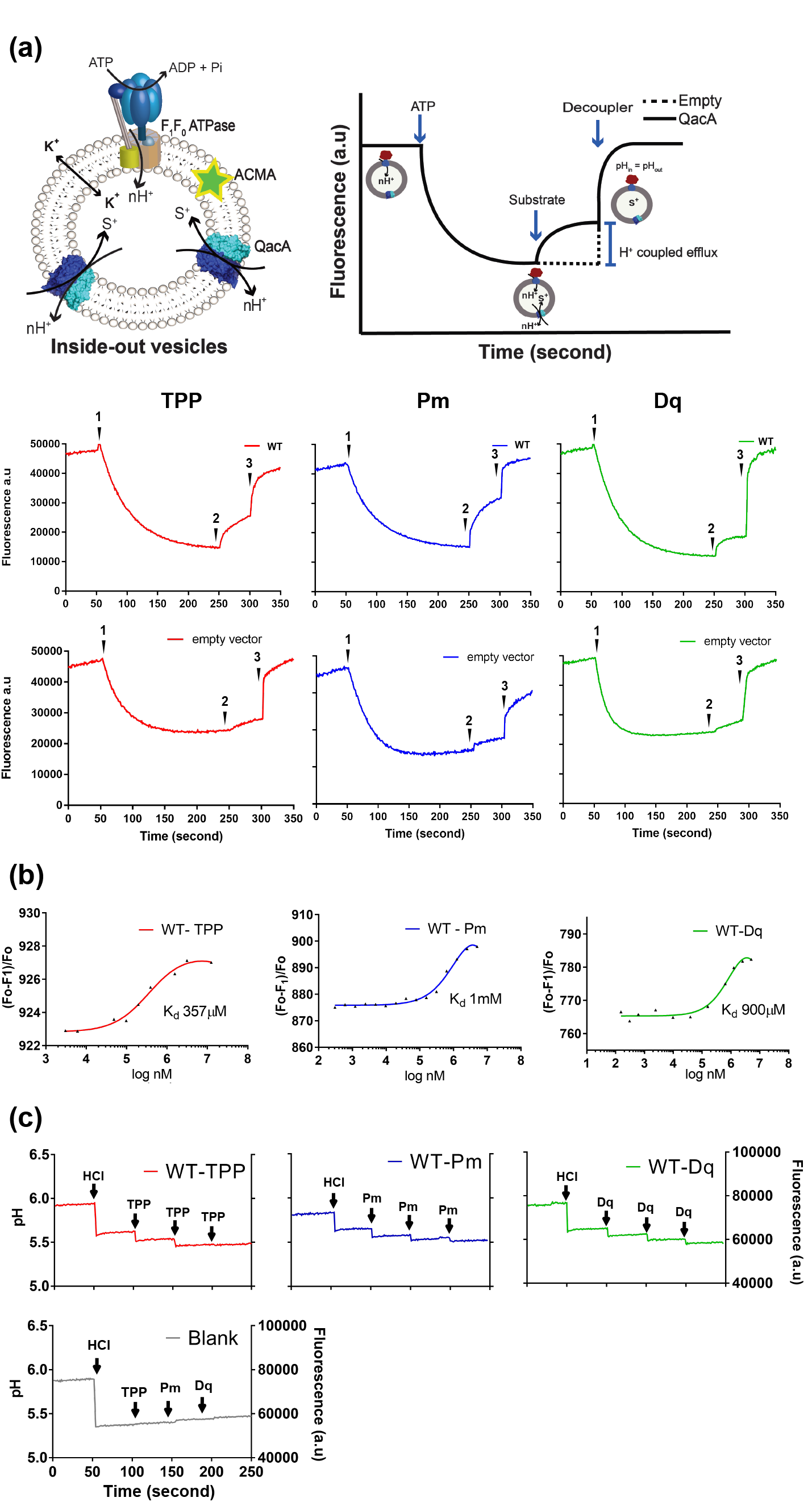
Functional characterization of WT QacA. (a) Schematic representation displaying substrate-induced H^+^ transport assay in inside-out vesicles (top panel); WT QacA is transport active for all the drugs tested (n=2). Addition of ATP is denoted by **1**; **2** represents substrate addition and **3** indicates addition of decoupler to abolish the H^+^ gradient. (b) The *K*_d_ estimated from the binding assays done using microscale thermophoresis displays higher μM to mM affinity for TPP, Pm and Dq (n=2). (c) Fluorescein’s fluorescence quenching experiment indicates a substrate-induced H+ release event. The assay is done with single batch of protein.

### Acidic residues in the vestibule exhibit variable importance to counter drug resistance

The alanine and asparagine/glutamine mutants at individual sites were tested alongside WT QacA for toxicity upon overexpression and none of the mutants displayed a toxic phenotype in the strain JD838, upon induction (Fig. S2a). A control mutation performed in the extracellular loop 7, D434N, had a significantly lowered expression (Fig. S2b). However, despite the minimal expression, no effect was observed in the drug resistance assays performed using Et, TPP, Pm and Dq, in comparison with WT QacA. FSEC was used to rapidly analyze expression levels and homogeneity of QacA and its mutants (Fig. 5). All the mutants exhibited detectable expression in the range of 44-125% in comparison to the WT (Table 1). The cells expressing these mutants were tested for their ability to grow at different dilutions using monovalent cationic substrates including Et, TPP and divalent cationic substrates including Pm and Dq (Fig. 6a). Experimentally optimized concentrations of substrates were used to perform the assay at neutral pH. QacA single mutants at D34 lost their ability to survive, irrespective of the cationic compound tested. This was followed by loss of activity with single mutants at D411 with most drugs, except Dq (Fig. 6b). Apart from D34, Dq had a loss of survival phenotype with E407 instead of D411, whereas the survival of other mutants remained unaffected in the presence of the drug. Also, Dq had no effect on the survival of D323 mutants despite being implicated in dicationic drug efflux, a phenomenon attributed to processive transport in an earlier study [35]. However, Pm had a pronounced effect on cell survival with minimal growth observed in most of the mutants including D323. The findings raise the interesting possibility of a few crucial sites essential for survival against antibacterial stress and distinct subsets of acidic residues required for drugs with different chemical structures.

**Figure 5.**
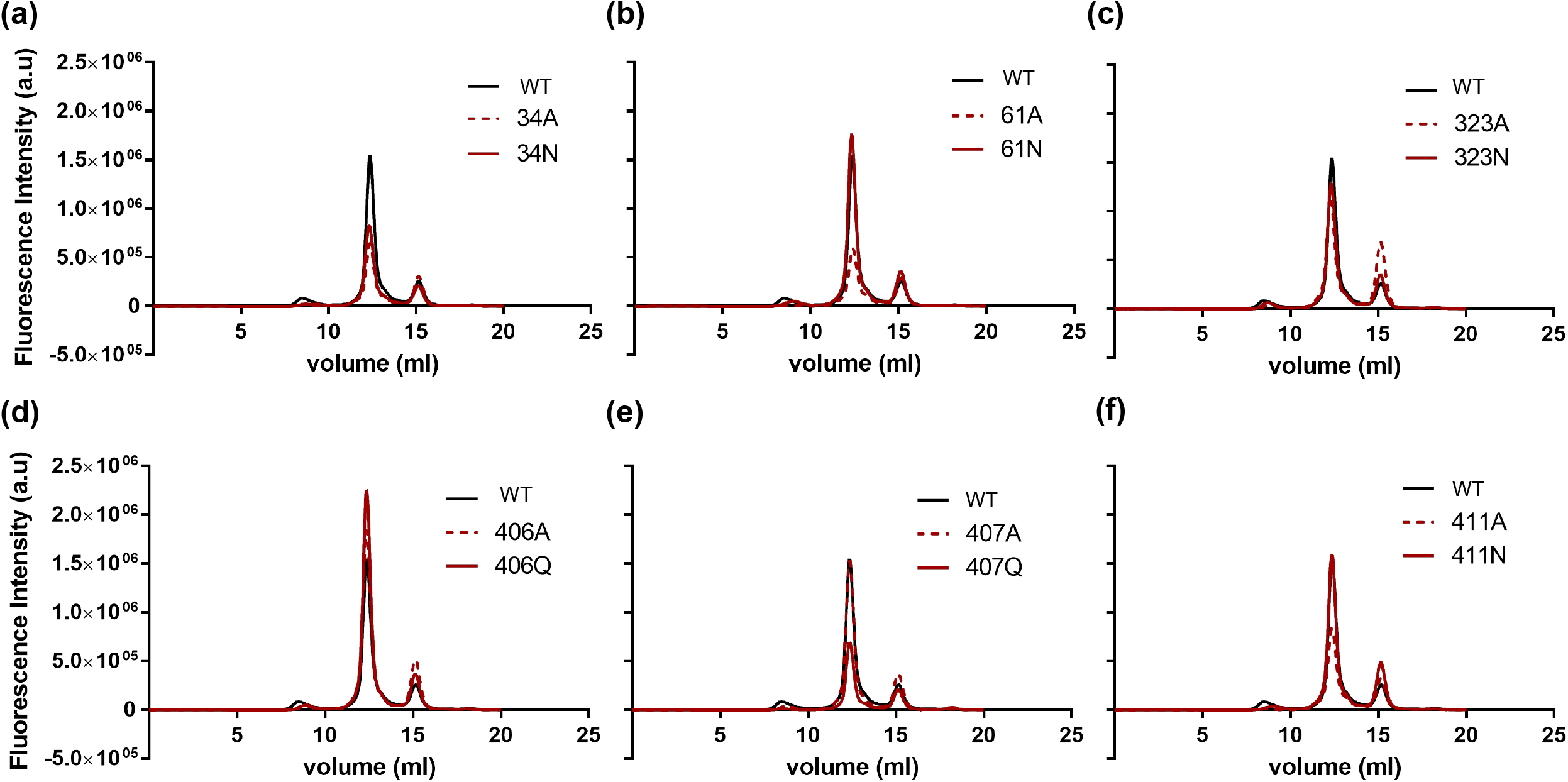
Fluorescence detection size exclusion chromatography (FSEC) traces (a-f) of WT QacA (black line) and its mutants. The Asn/Gln mutants (magenta) and Ala mutants (magenta-broken) along with WT QacA were expressed in *E. coli* with a GFP-His_8_ tag that allowed monitoring the expression profile in detergent solubilized crude membranes.

**Figure 6.**
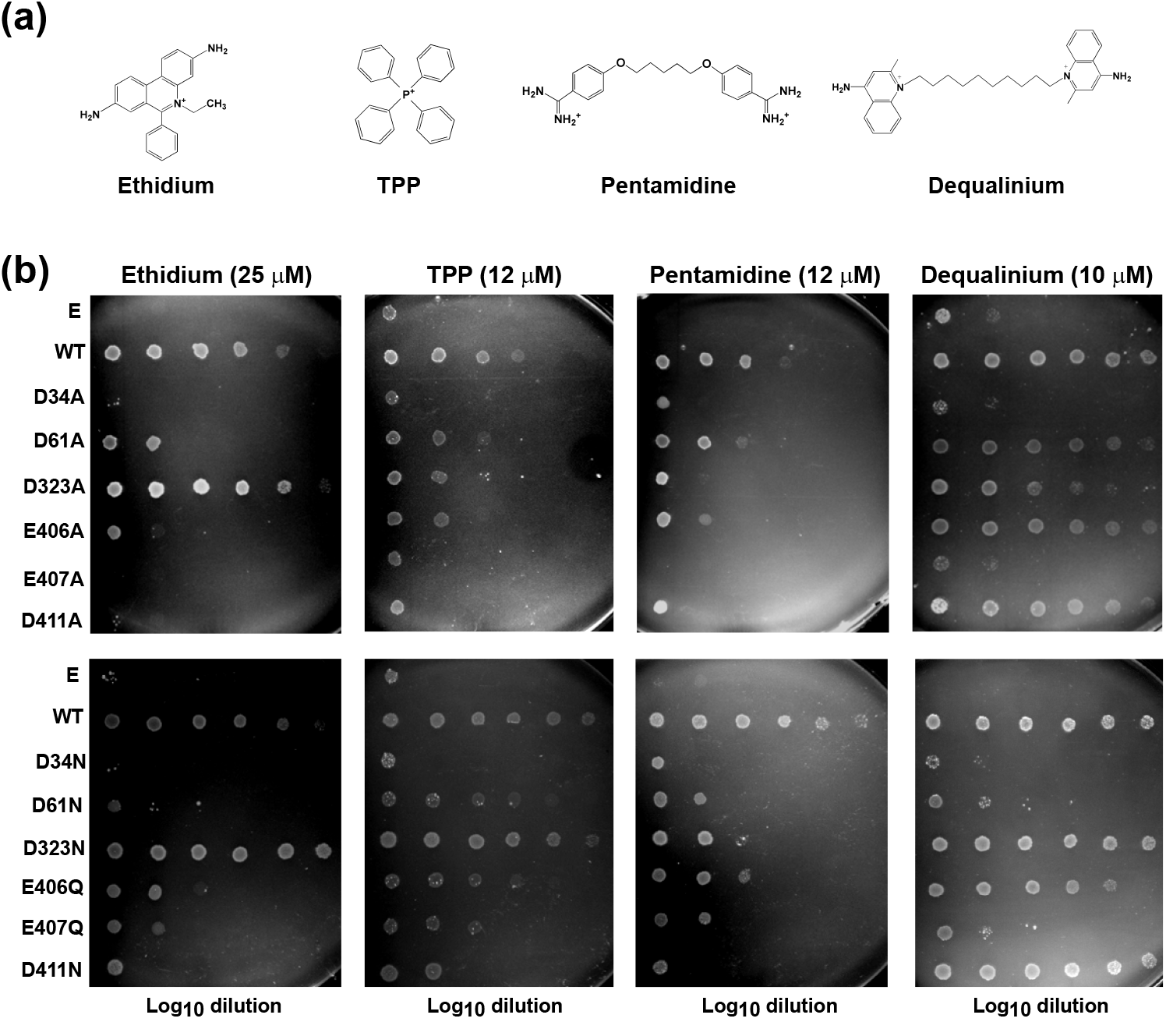
Drug resistance in *E.coli* cells expressing QacA. (a) Chemical structures of the cationic dye (Et) and antibacterial compounds (TPP, Pm, Dq) used as substrates for QacA. (b) Survival assay in presence of both monovalent (Et, TPP) and divalent (Pm, Dq) cations identifies four acidic residues D34, D323, E407 and D411 as crucial for the survival of bacteria in presence of toxic substrates. The role of D61 and E406 seems to be comparatively less significant. Log_10_ dilution starts with O.D. 1.0 at λ=600 nm from left to right. Plates are incubated for 12 hours (n=3).

**Table 1.** Area under curve was calculated to estimate relative expression levels of the mutants in comparison with WT QacA.

| Mutants | Area under curve<br>(in $10^6$) | % Relative<br>Expression |
| --- | --- | --- |
| WT QacA | 1.430 | 100 |
| D34A | 0.753 | 52 |
| D34N | 0.858 | 60 |
| D61A | 0.731 | 51 |
| D61N | 1.552 | 109 |
| D323A | 1.413 | 98 |
| D323N | 1.272 | 89 |
| E406A | 1.688 | 118 |
| E406Q | 1.781 | 125 |
| E407A | 1.346 | 94 |
| E407Q | 0.642 | 44 |
| D411A | 0.921 | 64 |
| D411N | 1.494 | 104 |

### Substrate recognition occurs at two sites in QacA

To analyze the roles of individual residues for substrate recognition and transport, we conducted whole cell-based Et efflux assays (Fig. 7), inside-out vesicle-based assays (Fig. 8, S3 and S4); substrate induced proton release assays (Fig. 9, S5) as well as affinity studies (Table 2, Fig. S6) in mutant backgrounds, with TPP, Pm and Dq. The alanine and isosteric substitutions at individual residues were analyzed for changes in transport properties of QacA. Isosteric substitutions of aspartate and glutamate are suggested to act as their permanently protonated versions [36], where the substrate cannot compete for the negative charge by displacing protons [36, 37]. A significant outcome of these assays is the characterization of D34 and D411 as essential sites for substrate recognition for most lipophilic cations used in this study (Fig. 7a, 7f, 8). D34 was predicted as analogous to E26 residue in the TM1 of MdfA [15] and D33 of VMAT[38], but its role was never investigated in earlier reports on QacA. Substitutions at D34 also compromise the ability of substrates to compete for protonation sites resulting in a near-complete loss of proton release as observed with TPP or retaining a minimal ability to compete for protons as observed in Pm and Dq (Fig. 9b). This loss of phenotype may stem from loss of measurable binding interactions with D34N QacA mutant as the dissociation constants could not be measured for any of the three drugs (Table 2, Fig S6).

**Figure 7.**
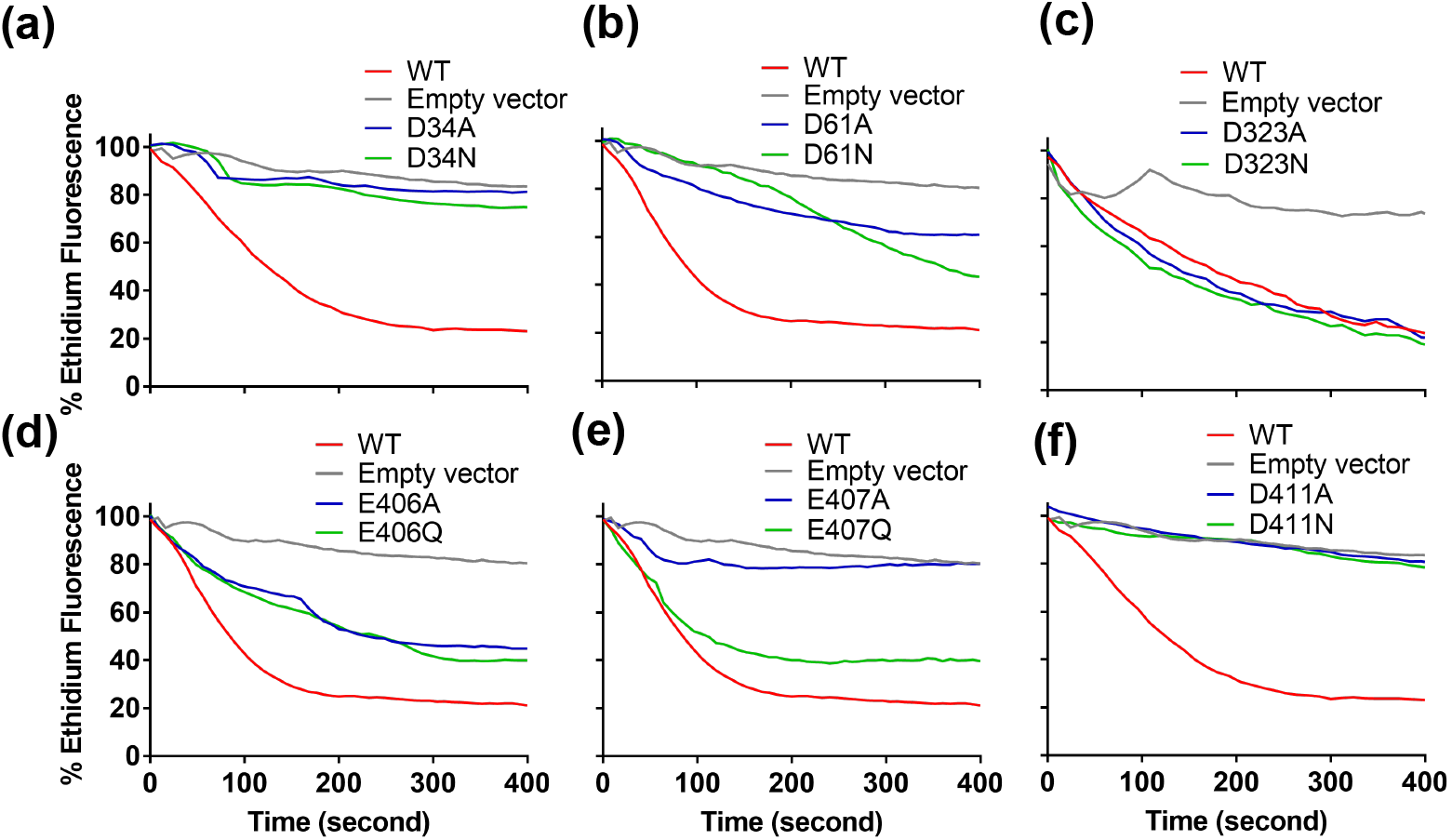
Altered transport properties of QacA mutants. (a-f) Whole cell efflux assay with WT QacA and its single mutants expressed in JD838 cells incubated with 50 μM EtBr for 1 hour in presence of CCCP (0.5 μM) (n=3). Altered efflux was measured as a function of ethidium fluorescence loss over a duration of 400 s.

**Figure 8.**
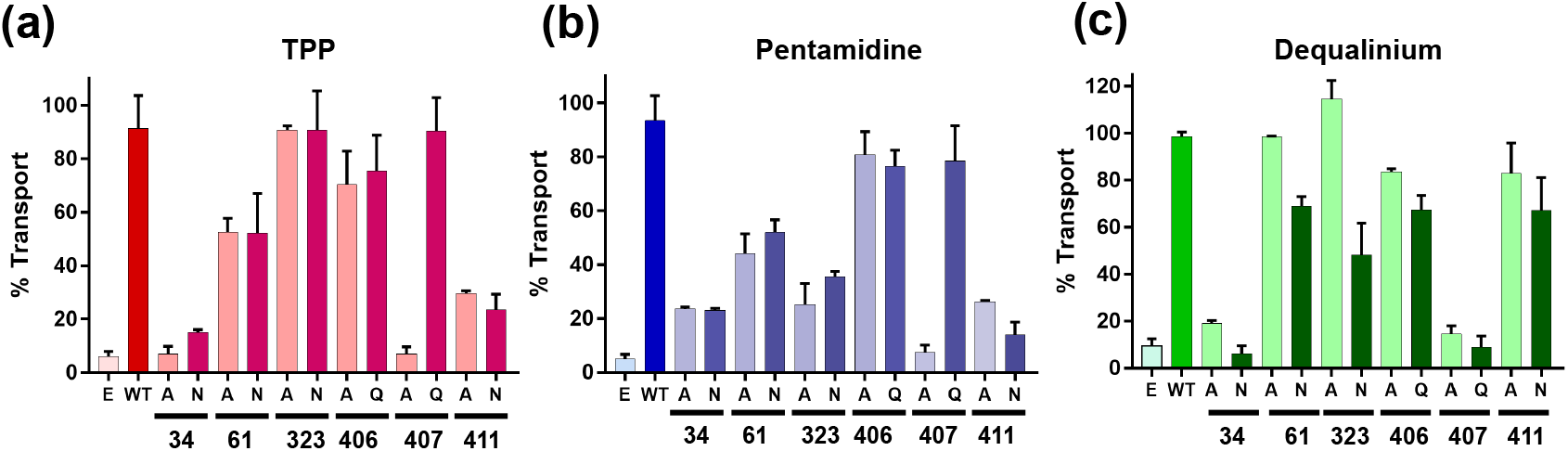
Inside-out vesicle based TPP, Pm and Dq induced proton transport (n = 2) assay. Valinomycin was kept throughout the experiment at 0.5 μM to prevent the build-up of membrane potential. The formation of ΔpH and the transport induced fluorescence dequenching of ACMA was measured (λ_Ex_= 409 nm, λ_Em_= 474 nm). The extent of ACMA flourescence dequenching, observed upon addition of the substrate to WT QacA vs. individual mutants is compared in the bar graph. Error bars represent the range observed for two independent measurements.

**Figure 9.**
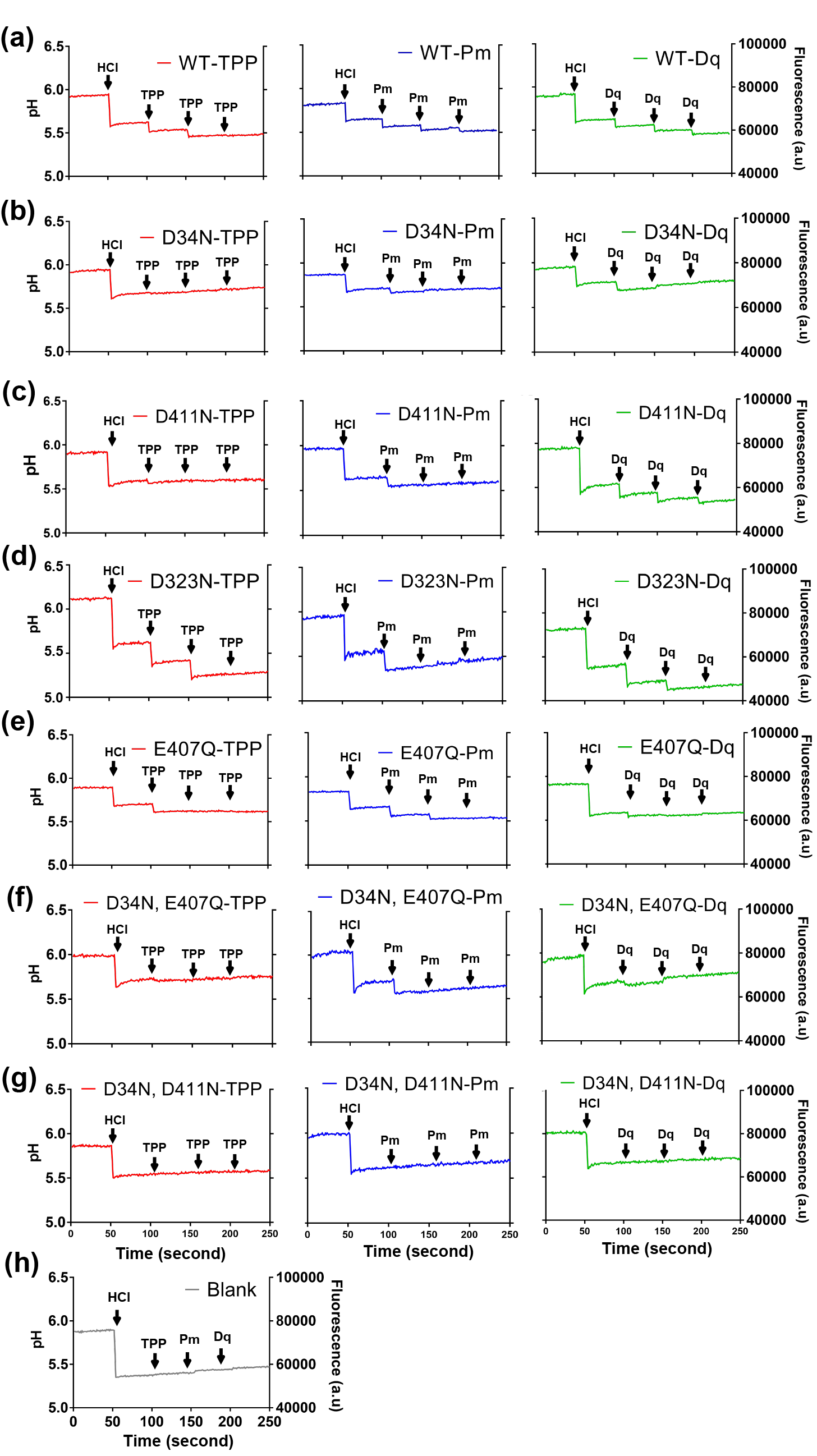
Substrate-induced proton release. The assay was done with purified QacA (6 μM, dialyzed to remove buffering agents). The pH of the solution was monitored in a time-dependent manner using fluorescein (2 μM), a pH-sensitive dye (λ_Ex_ = 494 nm, λ_Em_ = 521 nm). (a-g) Each row represents one mutant of QacA tested with three substrates. Arrows denote additions of HCl (6 μM) and titration of substrates (200 μM). (h) The titration was repeated in unbuffered solution without protein, as a negative control. A single batch of pure protein was used for these measurements.

**Table 2.** The dissociation constants (*K*_d_) are listed, analysed from the binding experiments done using MST (microscale thermophoresis) carried out with TPP, Pm and Dq.

| mutants | ligand | $K_d$ (mM) |
| --- | --- | --- |
| WT | TPP | $0.37 \pm 0.07$ |
| D34N | TPP | - |
| D61N | TPP | $0.94 \pm 0.10$ |
| D323N | TPP | $0.30 \pm 0.05$ |
| E406Q | TPP | $0.93 \pm 0.10$ |
| E407Q | TPP | $1.65 \pm 0.48$ |
| D411N | TPP | - |
| WT | Pm | $1 \pm 0.17$ |
| D34N | Pm | - |
| D61N | Pm | $0.96 \pm 0.20$ |
| D323N | Pm | - |
| E406Q | Pm | $1 \pm 0.13$ |
| E407Q | Pm | $0.98 \pm 0.19$ |
| D411N | Pm | - |
| WT | Dq | $0.90 \pm 0.21$ |
| D34N | Dq | - |
| D61N | Dq | $0.78 \pm 0.17$ |
| D323N | Dq | $0.50 \pm 0.10$ |
| E406Q | Dq | $0.87 \pm 0.10$ |
| E407Q | Dq | - |
| D411N | Dq | $0.71 \pm 0.04$ |

Interestingly, we found that D411 (TM13), topologically present towards the cytosolic end of the molecule, is also necessary for cells’ survival against all substrates, except Dq (Fig. 6). It was previously observed that D411C is accessible to PEG-Mal modification (Fig. 1c). Efflux experiments for Et, TPP and Pm indicate a crucial role of D411 in the activity of QacA, with both alanine and asparagine mutants displaying compromised activity for all three substrates (Fig. 7f and 8), whereas Dq’s ability to interact with QacA protonation sites remains unaltered in comparison to WT (Fig. 9c). The interaction propensities also reflect in the binding of TPP and Pm whose affinity to D411N could not be determined. However, Dq retains similar binding affinities with D411N and WT (0.71±0.04 mM vs 0.9±0.21 mM) (Table 2, Fig. S6), signifying the role of D411 as an important secondary substrate recognition site, specific to monovalent cations and divalent cationic substrates like Pm, but not Dq.

The presence of two distinct substrate recognition sites in QacA located in both cytosolic and extracellular halves of the transporter provides additional anchoring sites for the substrates to interact with, during the transport process. The finding that QacA employs different residues to recognize Dq as opposed to monovalent cations and Pm provides interesting hints into the promiscuity of substrate recognition that is explored in the subsequent section.

### D323 is required for pentamidine transport but not for other long-divalent cations

QacB, a paralog of QacA, has an alanine at residue 323 that was previously observed to impair transport of divalent cations like Pm and propamidine (Pi) [24]. However, recent observations suggest that it retains the ability to transport some divalent cationic drugs, particularly the ones separated by a long linker including Dq and chlorhexidine (Cx) [35]. As seen in the survival assays, QacA D323A/N mutants confer a stark contrast in the survival phenotype of the cells grown in the presence of monovalent cations vs Pm. While cells harboring D323A/N mutants survive well in presence of Et and TPP, the ability was lost in case of Pm, while survival in the presence of Dq remained unaltered (Fig. 6). Dq transport across inside-out vesicles remains active despite mutations at D323, indicating its importance specifically for Pm transport (Fig. 8b and c). A similar phenomenon was observed also with Cx where D323C mutation did not affect cell survival [35]. Also, with QacA D323N, the ability of the substrate to compete for protons in solution does not change with TPP and Dq in comparison with WT QacA (Fig. 9a and d). In line with the earlier experiments, Pm loses its ability to release protons from QacA D323N with a reduction in the number of H^+^- release steps, as compared to WT, prior to saturation. The importance of D323 for primarily interacting with Pm is also reflected in the binding affinities obtained for D323N, which binds to TPP and Dq with affinities comparable to WT QacA (*K*_d_ of ~0.3 mM for TPP and ~0.5 mM for Dq) whereas Pm shows a clear loss of affinity for the D323N mutant with accurate affinities not determinable due to limited solubility of Pm (Table 2, Fig. S6). The results indicate that D323 is not a crucial determinant of QacA’s ability to transport divalent cationic drugs but is important for specific substrates of QacA like Pm and likely Pi [24].

### E407 plays a dual role in substrate transport

E407 seems unique amongst the group of residues, as it is the only site whose loss of transport phenotype upon mutation to alanine is rescued by its isosteric mutation to glutamine. E407Q displays growth and efflux properties with Et, TPP and Pm comparable to WT QacA (Fig. 6, 7e, 8a and b) whereas E407A is incapable of transporting all three compounds. Interestingly, E407Q displayed a reduced number of titration steps by both TPP and Pm in comparison to WT QacA (Fig. 9e). Also, the binding properties of E407Q indicate a four-fold reduction in binding of TPP (*K*_d_ 1.65±0.5 mM) whereas Pm affinity remains unaltered (*K*_d_ 0.98±0.2 mM) in comparison to WT (TPP *K*_d_ 0.37±0.07 mM, Pm *K*_d_ 1±0.17 mM) (Table 2, Fig. S6), implying that it plays a minimal role as a substrate recognition site, even though it is needed for the survival of the host. Our observations indicate the likelihood that E407 acts as a protonation site in QacA given the ability of neutral substitutions of acidic residues to decouple transport from pH gradients [37]. Despite the proximity of E406 to E407, similar mutations at E406 do not elicit a major change in transport properties, suggesting that E407Q effects are indeed specific and cannot be substituted by neighboring residues.

While E407 serves as a potential protonation site for TPP and Pm (Fig. 10a and b), in the case of Dq, E407 is essential for substrate recognition (Fig. 10c). The survival assays in presence of Dq demonstrate that cells harboring E407A/Q mutants fail to grow whereas D411A/N mutations exhibit normal growth (Fig. 6). This apparent swap in their roles with respect to Dq strongly supports the promiscuity observed in the substrate repertoire of QacA, given that these residues are only one helical turn apart. In such a case, it would be plausible to argue that E407 acts as a substrate recognition site for one of the two charged moieties of Dq. This is supported by a range of observations, including rescue of phenotype observed in the background of E407Q mutant in the proton release driven by Et, TPP and Pm but failure of getting such observation in the same assays conducted for Dq efflux irrespective of A or Q substitution at E407 (Fig. 7e, 8). E407Q mutant has a similar affinity for Pm (*K*_d_ 1.00±0.17 mM in WT versus *K*_d_ 0.98±0.19 mM in E407Q) and reduced affinity for TPP (*K*_d_ 0.37±0.07 mM in WT vs. *K*_d_ 1.65±0.48 mM in E407Q), but is completely lost for Dq.

**Figure 10.**
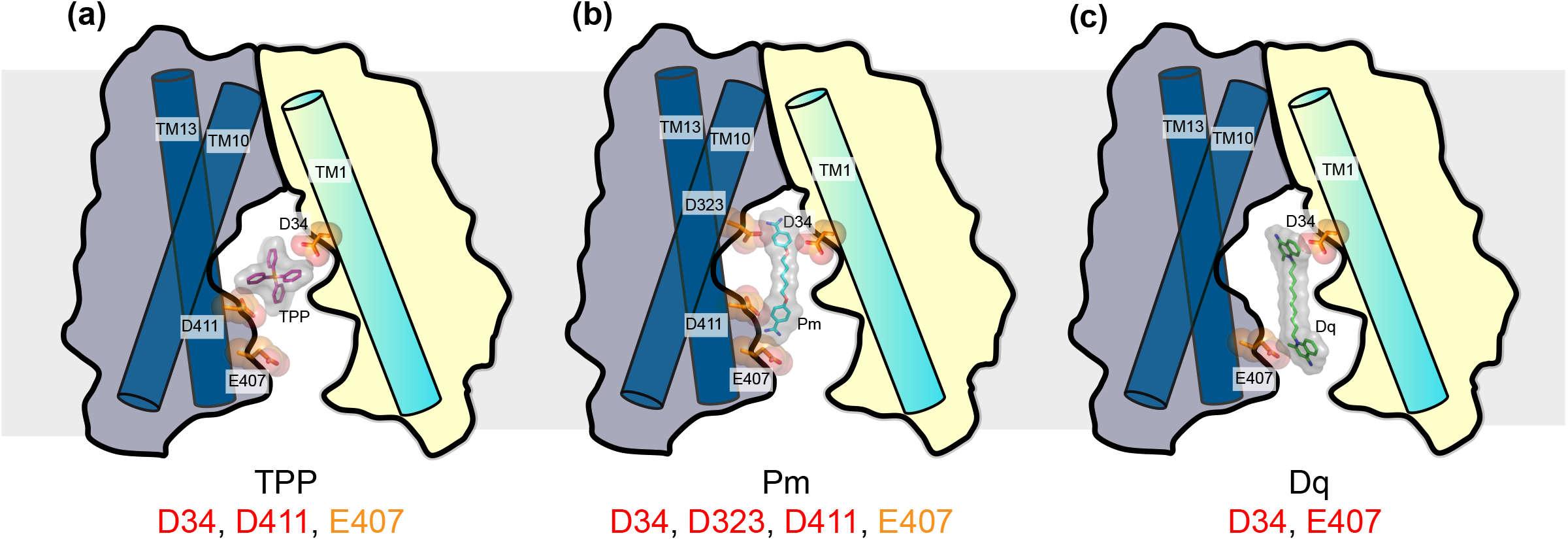
Substrate recognition and promiscuity. Cartoon depicting subsets of acidic residues that are important for individual cationic ligands. Residues identified as important for substrate recognition are colored in red and protonation sites helping in transport are colored in orange. (a) TPP transport requires D34, D411 and E407; (b) Pm, a biguanide requires four residues for transport; (c) Dq recognition and transport requires just two residues. The longer linker in Dq compared to Pm could result in the shift of the secondary recognition site from D411 to E407.

We made two double amino acid substitutions in QacA, namely D34N/E407Q and D34N/D411N to evaluate the effect on substrate-induced H^+^-release (Fig. 9f and g). In these combination mutants, TPP induced proton release was not observed with either of the two mutants since the D34N substitution alone was debilitating for TPP induced H^+^ release. Pm could induce limited proton release in D34N/E407N combination, attributing it to an unaltered D411. Dq, on the other hand did not seem to interact with any of the mutant combinations, likely owing to the important role played by D34 for the same.

The results strongly support the notion of discrete sets of residues being important for recognition and transport of antibacterial substrates of QacA. The essential residues are characterized by an “all-or-none” effect on substrate interaction and transport properties with single site mutations. These previously uncharacterized roles of acidic residues in QacA vestibule could be applicable to related DHA2 transporters, due to the strong conservation observed in the identified sites. The study also identifies a hitherto unknown region at the cytosolic part of TM13 as a hotspot for multi-substrate recognition and protonation driven transport, in DHA2 members.

## Discussion

Earlier studies on QacA attributed the differences in the transport properties between QacA and QacB paralogues to a single substitution D323A in TM10 [17]. It was observed that QacB was incapable of transporting divalent cationic drugs. However, in subsequent studies including this work, QacA mutants at D323 have been found to be capable of transporting both monovalent and some divalent cations. The ability to transport long divalent cations was attributed to processive transport in MdfA and was implicated as a mechanism in QacA D323C mutant’s ability to transport Dq and Cx [35]. However, Pm binding and transport are heavily compromised despite Pm being a long divalent cation. Also, closely related (> 40% identity) homologues of QacA do not have a conserved acidic residue at D323 position. These findings clearly indicate that residues critical for both monovalent and divalent cation recognition lie elsewhere in the TM helices.

Studies involving DHA1 members MdfA, LmrP and BbMAT have highlighted the important roles that protonatable acidic residues play in substrate recognition and H^+^:drug stoichiometry [12, 16, 39]. Asp and Glu residues in the binding pocket are protonated and can act as sites where cationic substrates can compete with and release protons during substrate translocation or serve as protonation sites, required for conformational transitions to facilitate transport and render the process electrogenic by increasing the H^+^:drug stoichiometry. Proton release through indirect competition was also observed in MdfA, wherein binding of TPP to a site that is distant from D34 induced the release of protons [40]. MdfA, in the recent past, was used to explore the importance of protonation sites [10, 15]. Repositioning of the acidic residue E26 from TM1 to TM10 (V335E) maintained the efflux properties of MdfA [10]. Also, the introduction of additional negative charges in the TM region at G354E, allowed conversion of electroneutral drug efflux in MdfA, to electrogenic efflux [39]. The G354E mutation in MdfA is localized in the cytosolic half of TM11 in MdfA that correlates to the position of E407 and D411 in TM13 in the QacA model.

Our findings on QacA suggest that it naturally encompasses several of the aforementioned findings observed in engineered DHA1 family antiporters. Its primary substrate recognition site D34 is akin to the substrate recognition sites of MdfA (E26), VMAT (D33) and BbMAT (D25) and is located in TM1. Although implicated in an earlier study using sequence alignments [41], we report the irreplaceable role of D34 for both monovalent and divalent cation recognition. Substitution to alanine or asparagine at D34 is not tolerated at this position irrespective of which cationic ligand is being transported. Although minimally accessible to PEG-Mal in solution (suggesting low accessibility to lipophilic cations), in induced H^+^ release experiments with D34N mutants, the substrates completely lose their ability to compete with protons, a likely indicator of indirect competition [40].

Locations for a secondary site for substrate binding is evident from our experiments at the cytosolic region of TM13, facing the vestibule. Among the three residues E406, E407 and D411, E406 is the least important for influencing transport. It is evident that D411A/N mutants’ transport properties towards Et, TPP and Pm were completely compromised, akin to the D34 mutants. TPP and Pm also had compromised binding in case of both D34 and D411 mutants, further reinforcing the importance of the two residues. However, E407A/Q mutants had rather dissimilar effects on transport properties. Interestingly, earlier studies with Cx and Pi have demonstrated reduced Et efflux through non-competitive inhibition [24]. In the light of our findings it is highly likely that non-competitive inhibition occurs primarily through interactions at the TM13 acidic residues.

While E407A mutant completely lost the ability to transport Et, TPP and Pm, E407Q retained WT QacA like transport properties. It is suggested in case of uniporters that acidic residue substitutions to neutral amino acids result in decoupling of substrate transport from pH gradients [36, 37]. The recovery of normal transport with E407Q suggests the residue’s role as a likely protonation site in QacA. E407Q also retained affinity towards most substrates used in this study. With Dq, E407 was observed to be essential for binding and transport, unlike Pm and TPP. Mutation at D411 exhibited near normal binding affinity for Dq. This selective difference in interactions based on the characteristics of the transported substrate is a testimony of QacA’s promiscuous substrate recognition properties (Fig. 10). The differences in site specificities for drugs like Pm and Dq could be due to variations in the length of the linker region. It is likely easier to accommodate the second cationic charge of Dq, which has a substantially longer linker compared to Pm, at E407 compared to D411, which is one helical turn above E407.

It has recently been proposed that MFS antiporters are capable of processive transport that likely involves a peristaltic movement of long dicationic substrates in two transport cycles instead of one, using MdfA and QacA D323C as test cases [35]. Although the phenomenon would require validation through structural snapshots of multiple intermediates in the transport cycle, our identification of secondary interaction sites towards the cytosolic half of QacA can greatly aid in understanding processive transport through simultaneous interactions with dicationic substrates, during the transport cycle. As mentioned earlier, the cluster of acidic residues identified in TM13 are well conserved in DHA2 members related to QacA (Fig. 1b). Future structural studies on DHA2 members can therefore be highly useful to understand and validate the roles played by protonatable residues in the vestibule of DHA2 members and identify strategies towards the design of specific efflux pump inhibitors against this class of drug efflux transporters.

## Acknowledgements

The authors would like to thank Prof. B. Gopal, Dr. Vinothkumar Kutti Ragunath and Dr. Abhijit A. Sardesai for critical reading of the manuscript. Authors are grateful to Ms. Swati Dubey, CDFD and Dr. Abhijit A. Sardesai, CDFD for the JD838 strain. PM is supported by the IISc-GATE PhD fellowship. AA is a student of the IISc-Integrated PhD program. NH is support by DBT-JRF PhD fellowship. AP is an intermediate fellow of the DBT-Wellcome Trust India Alliance (IA/1/15/2/502063) and a recipient of the Innovative Young Biotechnologist Award (IYBA) (BT/09/IYBA/2015/13) from the Dept. of Biotechnology (DBT), India.

## Author Contributions

PM and AP designed the research; PM, SK performed the experiments with help from NH and AG; AA and AP carried out homology modelling and analyses; PM, AA and AP analyzed the data and wrote the manuscript.

## Methods

### Plasmids and Strains

JD838 (∆*mdfA*∆*acrB*∆*ydhE*::Kan) strain is a derivative of the *E. coli* K-12 strain, LMG194, bearing a knockout of 3 multi-drug efflux pumps. A codon optimized QacA synthetic gene [34] was cloned into pBAD-His_8_ vector between *NdeI* and *HindIII* sites. CGFP construct was generated using megaprimer based whole plasmid PCR method. Site directed mutagenesis was performed using individual mutant primers and confirmed by DNA sequencing. The mutants of QacA include D34A/N, D61A/N, D323A/N, D411A/N, D434A/N, E406A/Q, E407A/Q, D34N/E407Q and D34N/D411N.

### Homology model and sequence analysis

Initially QacA ∆432-474 construct was modelled through homology modeling on I-TASSER server [31]. Known structures of homologous proteins and secondary structure restraints were used to thread the helices while a known structure of a POT transporter (PDB ID: 4IKV) was used to model QacA. One of the output structures which satisfied the structural characteristics of MFS transporter was chosen. The deleted region from residue no. 432-474 was built as a loop in the existing model using Completionist [42]. The model was embedded in a POPC bilayer made using CHARMM-GUI (http://www.charmm-gui.org) and was energy minimized using charmm36m forcefield [43]. To check the stability of the model, the whole system was simulated for 20 ns in explicit solvent using Gromacs 5.1.4 package [44] with a semiisotropic pcoupltype for equilibration and isotropic pcoupltype with Verlet scheme (md-vv) for production simulations. When the Cα RMSD stabilized, one frame was selected as the final model from the trajectory.

### Accessibility assay

WT QacA lacks cysteine residues in its sequence. Hence, the accessibility of acidic residues present in the vestibule to PEG-Mal was checked by generating cysteine mutants of D34, D61, D323, E406, E407 and D411 systematically. The purified proteins were incubated with PEG-Mal (0, 100, 250 μM) for 30 minutes at room temperature, followed by addition of 1 mM of β- mercaptoethanol for 5 minutes and incubation at room temperature to chemically inactivate excess PEG-Mal. The samples were loaded onto a 12% SDS-PAGE along with similarly treated but chemically unmodified WT QacA as a control. The experiment was repeated twice.

### Drug resistance assay

Resistance to the cationic compounds (Et, TPP, Pm and Dq) was assessed using JD838 cells expressing WT QacA and its alanine or isosteric mutants. For experiments on solid medium (2% w/v LB and 1.8% w/v agar), cells were diluted to an OD_600_ of 1.0, and 2 μl of a series of 10-fold dilutions were spotted on LB plates containing 0.05% (w/v) L-arabinose and 100 μg/ml ampicillin with or without the addition of the toxic cationic substrates (25 μM of Et, 12 μM of TPP, 12 μM of Pm and 10 μM of Dq). To check for the resistance against the toxic substance, the growth was analysed after 12 h incubation at 37°C. Similarly treated empty plasmid containing cells were used as a control. The assay was performed three times independently.

### Preparation of inside-out vesicle

JD838 cells expressing WT QacA or its mutants were grown for 12 h at 20^°^C in presence of 0.05% (w/v) L-arabinose and 100 μg/ml ampicillin. Cell were harvested and washed with 50 mM potassium phosphate buffer (pH 7) and resuspended in the same buffer containing 1 mg/ml of lysozyme. After 30 minutes of incubation at 30°C, cells were broken with 5-8 passes at ~500-600 bar pressure using a high-pressure homogenizer. The crude membranes were incubated for 30 minutes at 30°C with 10 mM MgSO_4_ and 10 μg/ml of DNase I. Unbroken cells and cell debris were removed by centrifugation at 13000g for 10 minutes at 4°C. Inside-out membrane vesicles were then isolated by ultracentrifugation at 125000g for 1h at 4°C and resuspended in 50 mM potassium phosphate buffer (pH 7) and 10% (v/v) glycerol. Vesicles were then frozen in liquid nitrogen and stored at -80°C for further use.

### Substrate-induced H^+^ transport in inside-out vesicles

Inside-out vesicles were prepared as mentioned above. The frozen vesicles were thawed quickly for 50 seconds at 46^°^C for the assay. Each measurement was done with 24 μg protein containing membranes diluted in a 2 ml solution of 50 mM KCl and 10 mM MgSO_4_. In the beginning of the experiment, 4 μM ACMA (9-amino-6-chloro-2-methoxyacridine) and 0.5 μM valinomycin were added and incubated for 5 minutes. The samples were continuously stirred during measurement of ACMA fluorescence (λ_Ex_=409 nm, λ_Em_=474 nm) using a Fluoromax-3 (Horiba) fluorescence spectrophotometer. ATP (100 μM) was added at 50^th^ second to create pH gradient required for transport process of QacA. As the gradient was established at 250^th^ second, 1 mM substrate (TPP, Pm or Dq) was added externally and the change in pH due to transport of the substrate was measured. The reaction was terminated using 4 μM nigericin at 300^th^ second. The assay was performed with two independent batches of inside-out vesicles.

### Whole cell efflux assay

For measuring Et efflux, JD838 cells were grown at 37°C for 3.5 h in presence of 100 μg/ml ampicillin and 0.05% (w/v) L-arabinose. After the cells were washed three times and resuspended in 20 mM HEPES (pH 7), 50 μM EtBr was added and incubated with for 1 h at 37°C in presence of 0.5 μM CCCP. Cells loaded with Et were washed three times with the same buffer. D-glucose (5 mM) was used to energize the cells. The efflux assay (λ_Ex_=530 nm and λ_Em_=610 nm) was done using Varioskan Flash (Thermo Scientific) plate reader with intermittent shaking. The assay was done in triplicate.

### QacA protein purification

Membrane preparation was done using standard protocol. The membrane was homogenized in 30 mM Phosphate buffer (pH 7) and 0.5 mM PMSF (Protease inhibitor) was added during homogenization. 20 mM UDM was used for protein extraction from the homogenized membrane by incubating with it for 1 h. Ultracentrifugation was done at ~125000g and the solubilized protein was present in the supernatant. Ni-NTA resin (pre-equilibrated in the buffer) was used for binding by nutating with it for 1 h at 4°C. The resin bound to protein was washed (with 30 mM Phosphate buffer (pH 7), 120 mM NaCl, 1 mM UDM, 30 mM imidazole) in a gravity flow column and eluted with 20 ml of elution buffer (300 mM imidazole, 30 mM phosphate buffer (pH 7), 120 mM NaCl, 1 mM UDM, 5% glycerol).. The protein was concentrated to 0.6-0.8 mg/ml prior to size exclusion chromatography (SEC). SEC was done with the purified protein using Superdex 200 increase column pre-equilibrated with 30 mM phosphate buffer (pH 7), 120 mM NaCl, 1 mM UDM and 5% glycerol.

### Determination of oligomeric status of QacA

SEC-Multi-Angle Light Scattering (MALS) was done using a three-detector system with light scattering (LS) detector, refractive index (RI) detector and UV detector to determine the molecular weight of QacA. The molecular weight of the protein was determined using protein conjugate mass analysis [45] with dn/dc = 0.18 for globular protein and dn/dc = 0.1506 for UDM. SEC-MALS (in 30 mM phosphate buffer (pH 7.0), 120 mM NaCl, 1 mM UDM and 5% glycerol (v/v)) was done with purified protein using Superdex 200 increase column.

### Binding assay

Binding assay was done using microscale thermophoresis (Nanotemper) [22]. Final protein concentration used for the assay was 50 nM. The protein was labeled with 50 nM red Tris-NTA dye in the C-terminus of the His-tag. Monolith^TM^ NT.115 MST premium coated capillaries were used in each experiment. In the binding reaction, the protein concentration was kept constant. The protein was incubated with 16 two-fold serial dilutions of the ligand. The ligand was solubilized in protein containing buffer (30 mM phosphate buffer, (pH 7.0), 120 mM NaCl, 1 mM UDM, 5% glycerol). The starting concentrations of the ligands were 10 mM in case of TPP and Dq, and 5 mM of Pm. The experiments were done with 2 independent sets of purified protein.

### Substrate-induced proton release assay

The assay was done with purified QacA (6 μM). Purified protein was dialysed with unbuffered solution (1 L; 200 mM NaCl, 0.8 mM UDM) to remove buffering agents. Fluorescein, a pH sensitive dye was used to detect the change in H^+^ concentration of the solution during the assay. Fluorescein (2 μM) was added at the start; at 50^th^ second 6 μM HCl and at 100^th^, 150^th^ and 200^th^ second 200 μM of the substrate (TPP, Pm and Dq) were added in the respective experiments. Control experiment was done in blank (200mM NaCl, 0.8 mM UDM) solution following the same procedure [35, 46]. Total duration of the experiment was 250 seconds. Fluorescein’s fluorescence was measured at λ_Ex_ = 494 nm, λ_Em_ = 521 nm in Fluoromax-3 (Horiba) fluorescence spectrophotometer with integration time of 1 second. The experiment was performed with a single batch of protein.

## Footnotes

**Authors declare no conflicting interests.**

**Email for correspondence**.; Email for strain requests.

## Abbreviations

ACMA: 9-Amino-6-Chloro-2-Methoxyacridine
TM: Transmembrane
DHA: Drug-Proton Antiporter/Drug-H^+^ Antiporter
Et: Ethidium
TPP: Tetraphenylphosphonium
Dq: Dequalinium
Pm: Pentamidine
Cx: Chlorhexidine
CCCP: Carbonyl cyanide m-chlorophenyl hydrazone
Pi: Propamidine
POT: Proton-dependent Oligopeptide Transporter
λ_Ex_/λ_Em_: Excitation/Emission Wavelength
MST: Microscale Thermophoresis
EL: Extracellular Loop
FSEC: Fluorescence-detection Size Exclusion Chromatography
UDM: Undecyl-β-D-Maltopyranoside
PEG-Mal: Polyethylene Glycol–Maleimide
MALS: Multi-Angle Light Scattering.

